# Droplet size and surface hydrophobicity enhance bacterial plasmid transfer rates in microscopic surface wetness

**DOI:** 10.1101/2022.03.07.483261

**Authors:** Tomer Orevi, Søren J. Sørensen, Nadav Kashtan

## Abstract

Conjugal plasmids constitute a major engine for horizontal gene transfer in bacteria, and are key drivers of the spread of antibiotic resistance, virulence, and metabolic functions. Bacteria in terrestrial habitats often inhabit surfaces that are not constantly water-saturated, where microscopic surface wetness (MSW), comprised of thin liquid films and microdroplets, permanently or intermittently occurs. How physical properties of microdroplets, and of the surfaces they reside on, affect plasmid transfer rates is not well understood. Here, building on microscopy-based microdroplet experiments, we examined the relation between droplet properties (size and spread) and plasmid transfer rates at single-cell and individual droplet resolution, using *Pseudomonas putida* as a model species. We show that transfer rates increase with droplet size, due to higher densities of cells on the surface in larger droplets, resulting from lower ratio between the area of the liquid-solid interface and droplet volumes. We further show that surface hydrophobicity promotes transfer rates via the same mechanism. Our results provide new insights into how physical properties of surfaces and MSW affect plasmid transfer rates, and more generally, interactions mediated by cell-to-cell contact, with important implications for our understanding of the ecology and evolution of bacteria in unsaturated environments.

## Main

Conjugative plasmids confer countless important bacterial traits, including antibiotic resistance and pathogenicity, and are a major vehicle of horizontal gene transfer within microbial communities [1-5]. In terrestrial ecosystems, bacteria commonly inhabit surfaces that are not constantly saturated with water, which are often covered by thin liquid films and microdroplets, termed microscopic surface wetness (MSW) [6-8]. Under MSW conditions, cells are confined within droplets or films for prolonged periods, and thus cell-to-cell interactions mediated by physical contact, which is required for conjugation, are expected to be high. Conjugation also depends upon the cells’ physiological state [9-11], which is in turn affected by the unique physicochemical conditions of MSW [6]. A number of studies have elucidated the mechanisms that modulate plasmid transfer rates on surfaces and in porous media at the microscale [12-15], as well as the various ways in which hydration conditions and dynamics affect conjugation rates [10, 12, 16, 17]. Yet, how basic physical properties of individual microdroplets (e.g., their dimensions) and those of the surfaces upon which they reside, affect plasmid transfer rates has not been studied in-depth.

Here, we use a microscopy-based microdroplet experimental system wherein we spray media-suspended bacterial cells onto a surface, enabling us to track plasmid transfers within individual microdroplets (see Methods in SI). We use strains of *Pseudomonas putida* KT2440 [18] – a well-studied model bacterium for unsaturated environments – as plasmid donors and recipients. The donor strain carries a conjugal plasmid with a GFP reporter that is repressed in the donor cells but expressed in trans-conjugant cells [4].

We first asked how droplet size affects the number of plasmid transfer events (T_e_) within individual droplets. We sprayed bacteria suspended in liquid M9 minimal media onto glass-bottom 12-well plates. Then, plates were kept at 28°C and at relative humidity (RH) of ≈98% (SI). We captured microscopy images of the wells’ bottom surfaces at 0, 3, 6, 12, and 18 hrs post inoculation. Under these conditions, evaporation rate was slow and droplets did not dry out, while cells within them grew very slowly (Table S1). The area of the liquid-solid interface of each sessile droplet (the contact area between the droplet and the surface, which we term ‘droplet area’was used as a quantitative measure of droplet size. Droplet area spanned several orders of magnitude ranging from ∼10^2^ µm^2^ to ∼10^5^ µm^2^ (Fig. 1A,B).

**Fig. 1.**
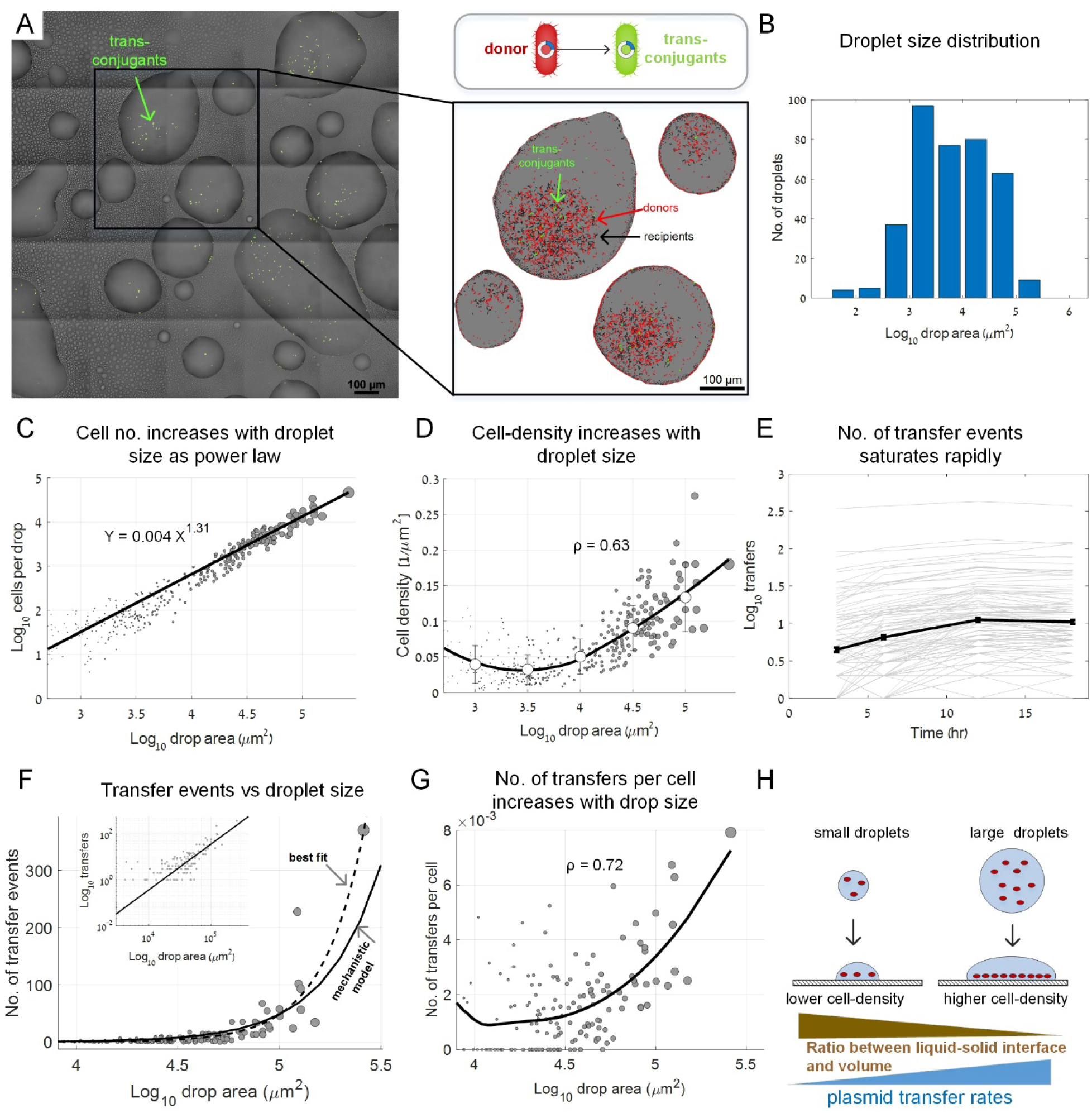
Droplet size effect on plasmid transfer rates. **A**. A representative section of the glass surface covered by sprayed droplets at 6 hrs after spraying. Green cells are trans-conjugants. Inset: zooming into individual droplets. Donor, recipient, and trans-conjugant cells are shown in red, black, and green respectively. **B**. Droplet area histogram (‘area’ refers to the area of the liquid-solid interface, A). **C**. The total number of cells (N) in individual droplets (including donors, recipients, and trans-conjugates at t = 6 hrs). Circle size reflects droplet area. Axis plotted in log-log scale (dataset includes 372 droplets). **D**. Surface cell density in individual droplets. Line shows smoothed average (using LOESS) circles represent mean±STD for binned data. ρ represents Spearman rank correlation coefficients (P < 10e-10) **E**. Lines show cumulative no. of transfer events over time in individual droplets at 3, 6, 12, and 18 hrs post inoculation (semi-log scale). Solid black line represents mean±SE (SE smaller than symbols). **F**. No. of transfer events as a function of droplet area: Circles are experimental data. Black line: mechanistic model T_e_ = 3.58·10^−7^·A^1.62^ (SI). Dashed line: best-fitted line based on data T_e_ = 1.09·10^−7^·A^2.12^ (fitted as a power law, SI). Inset: same data but at log-log scale. **G**. Transfer events per cell (T_c_) as a function of droplet area. Black line is smoothed data (LOESS). ρ represents Spearman rank correlation coefficients (P < 10e-10). **H**. Conceptual representation of the underlying mechanisms explaining the increase in plasmid transfers as a function of droplet area.

We focused our analyses on the liquid-solid interface where we observed that the majority of cells settle and where most plasmid transfer events occurred. The number of cells (N) observed on the surface in each droplet increased as a power function of droplet area (A) with a slope >1 (N = 0.004·A^1.31^; Fig. 1C), reflecting a lower ratio between A and droplet volume in larger droplets. Importantly, though initially slightly decreasing, surface cell density increased with droplet area for droplets > 3,000 µm^2^ (Fig. 1D). Assuming equal cell densities in the sprayed droplets prior to surface deposition, in larger droplets a higher number of cells are deposited on a smaller area of the surface, leading to increased cell density (Fig. 1D). We hypothesized that the increased density in larger droplets would result in a higher number of per-cell transfers (T_c_).

We first tracked the number of transfer events (T_e_) within individual droplets. Trans-conjugates were observed as early as few hours post inoculation. At 6 hrs, there were already dozens of transfer events in the larger droplets (Fig. 1A,E). The majority of transfers occurred within 6 to12 hrs, and peaked at 12 hrs (Fig. 1E), corroborating other studies [10, 16, 17]. We decided to focus on the t = 6 hrs point for further in-depth analysis (see also Fig. S1).

Next, we sought to test whether T_e_ is indeed governed by cell density. Theoretically, in each droplet, the number of donor and recipient cells per unit area dictate the probability of donor-recipient physical contact (that is required for conjugation). As cells did not always remain in an exact location throughout the experiment, instead of counting donor-recipient pairs exhibiting physical contact, we employed a statistical approach utilizing data from hundreds of individual droplets for fitting a simple per-droplet, population-based mechanistic model [1, 19]: T_e_ = k · (D_d_ · D_r_ · A) (where D_d_ – donor density; D_r_ – recipient density; A – drop area; and k is a constant, SI). Indeed, analysis of a subset of droplets, wherein we counted all donor-recipient pairs within a 5 µm distance, showed that such a population-based approach can serve as a good approximation for the number of donor-recipient pairs that are located in close proximity (Fig. S2).

To develop a mechanistic model for T_e_ as a function of droplet area (A), we approximated D_d_ and D_r_ based on the empirical power-law relation (Fig. 1C) between the expected number of cells (N) and A, while taking into account the overall ratio of donor-to-recipient cells (1:3). Then we fitted the overall mechanistic model (by fitting k) to the empirical T_e_ data in individual droplets (Fig. S3, SI). This yielded the power function T_e_ = 3.58·10^−7^·A^1.62^ (see SI), which agrees well with the experimental data (Fig. 1F, see also inset). A best-fitted power-law model, that was not based on our mechanistic model, showed a somewhat higher exponent: T_e_ = 1.08·10^−9^·A^2.12^ (Fig. 1F, SI), possibly pointing to an additional positive effect of A on T_e_, or a higher ‘effective’ density within larger droplets (as can be seen in Fig. 1A inset, possibly due to droplet growth by condensation). We moreover found that not only does T_e_ increase with A, but so does the mean number of transfers per cell T_c_ (Fig. 1G). These results highlight the apparent underlying mechanism: a decrease in the ratio between the area the droplet covers on the surface and the droplet volume, leading to an increase in surface cell density with droplet size (Fig. 1H).

As the ratio between the area that the drop captures on the surface to its volume appears to be a key variable of the system, we next wanted to test how surface hydrophobicity affects the system. We hypothesized that a more hydrophobic surface would lead to a higher T_c_ due to a lower ratio between A and droplet volume that results in higher cell densities. To test this hypothesis, we modified the hydrophobicity of our glass-bottom well-plates to produce a hydrophobic or a hydrophilic surface (contact angles of ≈90° and ≈42° respectively, see SI and Fig. S4). We then performed similar microdroplet experiments with the modified surfaces (due to technical reasons we also changed additional settings of the original experiment, SI).

The average area of the sprayed droplets on the hydrophilic surface was indeed about twice that of the hydrophobic one (Fig. 2A,B) as reflected in the droplet area distributions (Fig. 2C, Fig. S5). The number of cells per droplet reflected that as well: There were more cells per given droplet area on the hydrophobic surface (Fig. 2D). Once again, cell density increased with droplet area, with higher densities on the hydrophobic surface (Fig. 2E).

**Fig. 2.**
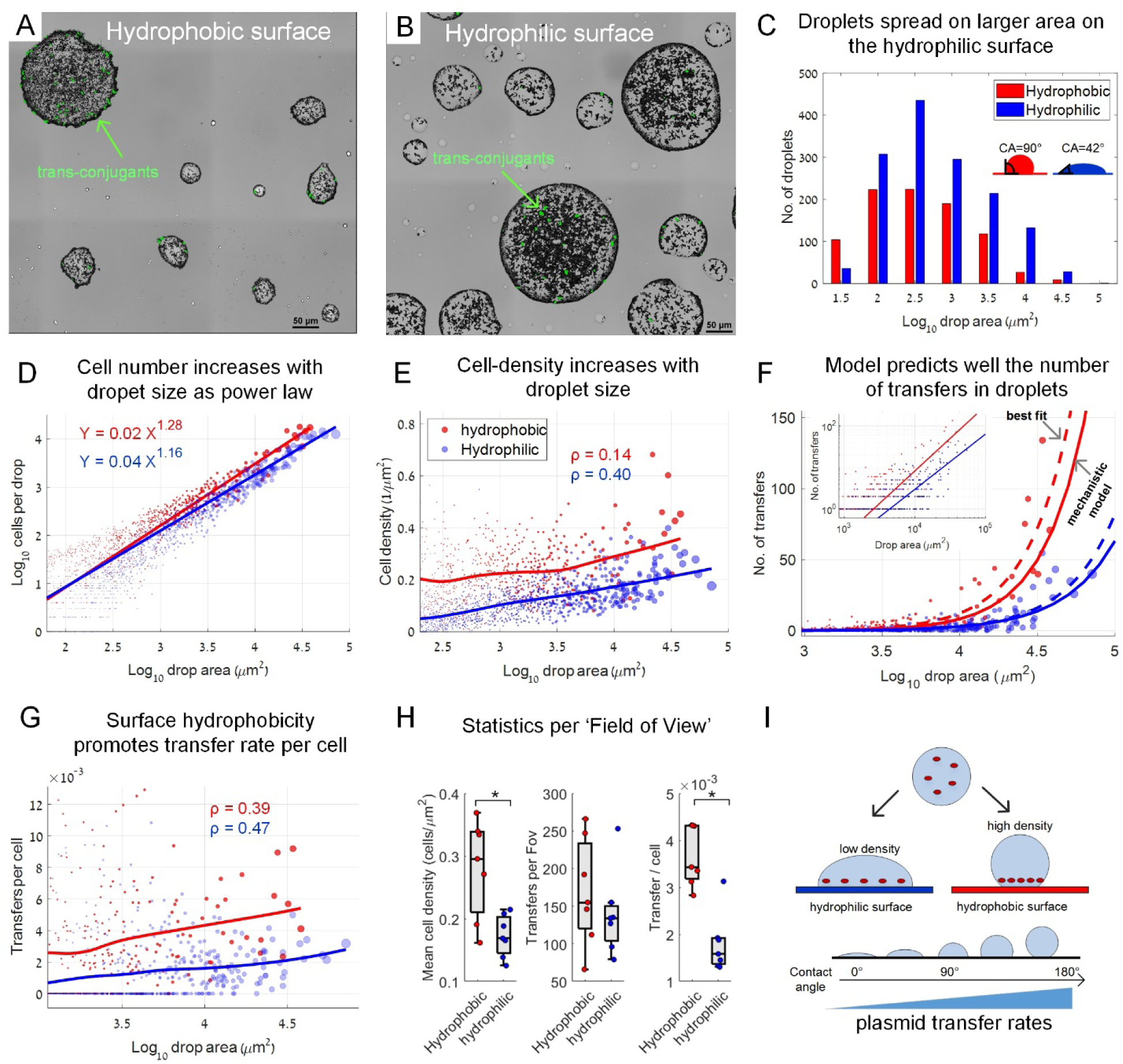
Surface hydrophobicity effect on plasmid transfer rates. **A**. A representative section of a hydrophobic surface covered by sprayed droplets at 6 hrs after spraying. Green cells are trans-conjugants. **B**. Same as in (A) but on a hydrophilic surface. **C**. Droplet area histograms (area refers to area of the liquid-solid interface). Contact angles (CA) were measured as described in SI. **D**. Cell numbers in individual droplets including donors, recipients, and trans-conjugates. Circle size reflects droplet area. Axis plotted on a log-log scale (dataset includes 2,518 droplets) **E**. Surface cell density in individual droplets. Line represents average smoothing (LOESS). ρ represents Spearman rank correlation coefficients (P = 0.0294 for the hydrophobic experiments, and P < 10e-10 for the hydrophilic one; ρ calculated for drops with A > 10^3^ µm^2^). **F**. No. of transfer events (T_e_) as a function of droplet area: Circles represent experimental data. Solid lines: mechanistic model (see SI). Dashed line: best-fitted line based on data (fitted as a power function, SI). Inset: same data but at log-log scale. **G**. Transfer events per cell (T_c_) as a function of droplet area. Solid lines represent average smoothing (LOESS). ρ represents Spearman rank correlation coefficients (P = 3.44e-10 for the hydrophobic experiment, and P < 10e-10 for the hydrophilic one; ρ calculated for drops with A > 10^3^ µm^2^). **H**. Statistics per entire surface sections (each of 1.6 mm x 1.6 mm). Boxplot represents median and 25%, 75% percentiles. Statistical comparisons were based on two-sample t-test yielding P = 0.0058, P = 0.41, and P = 1.34e-04 for the three comparisons respectively. * indicates statistical significance at 0.01 significance level. **I**. Conceptual representation of the underlying mechanisms explaining the increase in plasmid transfer rates as a function of surface hydrophobicity.

T_e_ increased with A for both surface types, concurring with our previous experiment (Fig. 2F). We found that the mechanistic model, with a fitted k (Fig. S6) and incorporating the cell:area ratio (Fig. 2D), agreed well with the observed T_e_ data (Fig. 2F, SI). A best-fitted model based on data only again showed a slightly higher increase in T_e_ as a function of A (Fig. 2F, SI). As before, T_c_ increased with A and was higher for the hydrophobic surface, as anticipated (Fig. 2G). The described trends are also valid for a cumulative analysis of entire surface sections that were scanned (Fig. 2H). In summary, surface hydrophobicity promotes T_e_ through increasing surface cell densities (Fig. 2I).

To summarize, our results provide new insights into the ways physical properties of surfaces and MSW affect conjugal plasmid transfer rates. We show that transfer rates are governed by cell density, which is dictated by droplet area and surface hydrophobicity. Our results further demonstrate how fluid mechanics and microscale wetting properties affect plasmid transfer rates, and more broadly affect interactions that require cell-to-cell physical contact. We note that higher bacterial densities in larger droplets are likewise expected due to other mechanisms, including favorable growth conditions and higher survival in larger droplets [7] and inter-species synergistic interactions due to higher diversity in larger droplets [20], suggesting that in real-world conditions, this link might be even stronger. As conjugative plasmids play a central role in horizontal gene transfer, the means by which the physical properties of the system affect plasmid transfer rates are important. This study has implications for the ecology and evolution of bacteria in unsaturated environments including the built environment, soil, root and leaf surfaces [8], and animal and human skin, with potential applications such as containing the management of the spread of antibiotic resistance.

## Acknowledgements

We thank Jonathan Friedman for valuable comments and discussions, and Roni Wallach and Felix Ogunmokun for their assistance with contact angle measurements. NK is supported by research grants from the James S. McDonnell Foundation (Studying Complex Systems Scholar Award, Grant #220020475) and the Israel Science Foundation (ISF #1396/19). SJS is partly supported by Grants NNF200C0062223 and NNF20OC0064822 from the Novo Nordic Foundation.

## Notes

### Competing Interest Statement

The authors have declared no competing interest.

