## Supplementary Information for "Droplet size and surface hydrophobicity enhance bacterial plasmid transfer rates in microscopic surface wetness"

**Orevi et al.**

SI Includes:

**1. Supporting Materials and Methods**

**2. Supporting Figures**

**3. Supporting Tables**

### 1. Supporting Materials and Methods

#### Bacterial strains and growth conditions

*Pseudomonas putida* KT2440 cells were used as donor and recipient pair. Donor cells were constructed by Sørensen & Smets as described elsewhere [1]. Briefly, donor cells are chromosomally tagged with constitutively expressed mCherry and lacIq production (and kanamycin resistance gene). Additionally, donor cells carry a broad host pKJK5 plasmid marked with gfpmut3b gene which is expressed by a LacIq repressible promoter Plac (plasmid includes kanamycin and tetracycline resistance genes). *P. putida* KT2440 cells were routinely cultured in M9 medium (M9 Minimal Salts Base 5x, Formedium, UK) supplemented with 20 mM Glucose and 50 µg/mL Km + 10 µg/mL tetracycline (for the donor strain), under agitation set at 220 rpm, at 28°C.

#### Experiment 1: Plasmid transfer in droplets sprayed on untreated glass substrate

Donor and recipient strain were cultured (separately) in 50 ml Falcon tubes continuing 25 mL of medium and appropriate antibiotic. Overnight cultures were washed twice (centrifuge at 6000 rcf for 5 min) in M9 medium and the pellets were re-suspended in 2 mL medium in order to reach high density cultures ($OD_{600}$ > 5). Next, the OD of the donor and recipient strains were adjusted to $OD_{600}$ = 4 and the strains were mixed with a 1:3 (donor:recipient) ratio in a 5 mL tube that continued 1 µM of Alexa dye (Alexa Fluor 647, Invitrogen) that was used to fluorescently stain the sprayed droplets. The solution was loaded into 5 mL refillable spray bottles (purchased at a local cosmetics store) and a portion of the load was sprayed on a 12-well glass bottom plate (P12-1.5H-N, Cellvis) in the following manner: a 12 well plate was placed (without the plastic lid cover) in a plastic bag, and then the solution was delivered by pressing the spray nozzle 4 times from a distance of about 15 cm above the plate. Tap water were added to the empty spaces between the wells of the plate, plates were covered with the plastic lid, and the plate's perimeter was sealed with a stretchable sealing tape to maintain a humid environment (>98% RH). The plate was incubated in the dark at $28^0$C throughout the duration of the experiment (18 hr).

**Experiment 2: Plasmid transfer in droplets sprayed on hydrophobic and hydrophilic modified glass substrate**

Experiment II was done similarly to experiment I with the following modifications: (a) The donor and recipient mix solution was adjusted to $OD_{600} = 3$ with a 1:1 ratio (b) spray was applied to each well separately through a cylinder made out of 50 mL Falcon tube from which the conical end was chopped 1.5 cm above the base of the tube (one press of the spray nozzle for each well) (c) hydrophobic and hydrophilic modified glass well plates were used.

**Glass modification**

Procedure for hydrophobic and hydrophilic glass modification was adopted from [2] with few adjustments: 12-well glass bottom plate (P12-1.5H-N, Cellvis) were filled with 2% RBS35 solution (RBS™ 35 solution, Sigma) and sonicated for 5 min in a ultrasonic bath (model ACP-200H, MRC) followed by thorough rinsing with tap water, demineralized water, methanol, tap water and finally demineralized water again to obtain a hydrophilic surface. The plates were dried in an oven for 2 hr in $70^0$C and either stored before use or further modified by applying a hydrophobic coating. To obtain a hydrophobic surface, 20 µL of 2% (v/v) dichlorodimethylsilane (CAS 75-78-5, Sigma) in trichloroethylene (CAS No 79-01-6, Sigma) were applied to the center of each well (avoiding contact between the siliconizing solution and the boundaries of the well which result in melting of the well plate coting) and left to dry for 1 hr in a chemical hood. The plates were then dried in an oven for 2 hr at $70^0$C, rinsed with tap water, demineralized water and dried in the hood for 2 hr before use.

**Microscopy**

On the indicated time points (see Main Text), 12-well plates were mounted on a stage top chamber (H301-K-FRAME, okolab) set at 28 °C. Microscopic inspection and image acquisition were performed using an Eclipse Ti-E inverted microscope (Nikon) equipped with Plan Apo 40x/095 N.A. air objective and the Perfect Focus System for maintenance of focus. A LED light source (SOLA SE II, Lumencor) was used for fluorescence excitation. GFP fluorescence was excited with a 470/40 filter, and emission was collected with a T495lpxr dichroic mirror and a 525/50 filter. mCherry fluorescence was excited with a 545/25 filter, and emission was collected with a T565lpxr dichroic mirror and a 605/70 filter. Alexa 647 fluorescence was excited with a 620/60 filter, and emission was collected with a T660lpxr dichroic mirror and a 700/75 filter. Filters and dichroic mirror were purchased from Chroma, USA. A motorized encoded scanning stage (Märzhäuser Wetzlar, DE) was used to collect multiple positions of the well bottom surface. In each well, two random positions were picked and imaged by scanning 5 × 5 adjacent fields of view (with a 5% overlap, 1.6 × 1.6 mm per scan). Images were acquired with an SCMOS camera (ZYLA 19 4.2PLUS, Andor, Oxford Instruments, UK). NIS Elements 5.02 software was used for acquisition.

### Image processing

Image processing and analyses were performed to quantify the area of the liquid-surface interface (droplet area), total number of cells per droplet, number of donor cells per droplet and plasmid transfer events per droplet. NIS Elements 5.02 software was used for image processing. droplet masks were generated by intensity threshold segmentation of the Alexa 647 channel. Binary masks were converted to region of interest (ROI) elements delineating droplets boundaries. Bright field channel was used to identify the total cell (donor, recipient and transconjugate) entities within single droplets (i.e., identified ROI's). Rolling ball background correction (0.49 µm) was applied on the entire image and the 'spot detection tool' was applied to enumerate cell number. mCherry channel was used to identify the donor cell entities within single droplets. Rolling ball background correction (0.49 µm) was applied on the entire image and 'spot detection tool' was applied to enumerate donor cell number. GFP channel was used to identify trans-conjugant cells within single droplets. Rolling ball background correction (0.49 µm) was applied on the entire image and intensity threshold was applied to identify single GFP expressing cells. The 'dilate' tool was operated on the resulting binary mask in order to cluster adjacent GFP expressing cells into a single object counted as a single 'plasmid transfer event'_ (i.e. we assumed the plasmid was acquired prior to cell division). See also Fig. S7.

### Data and statistical analysis

Data and statistical analysis were done with MATLAB version 2021b.

Experiment 1 (untreated glass surface): Dataset consisted of 12 surface sections of 1.6 mm × 1.6 mm (FoVs) from 6 different wells, with a total of 372 droplets. Power function coefficients were estimated using 'fitnlm' function to a simple power function of the form: $Y=bX^a$. Data smoothing were done using 'smooth'(X,Y,r,'loess') with r=0.8 in Fig. 1D and r=0.5 in Fig 1G. Spearman rank correlation coefficients in Fig.1D: rho=0.63 P< 10e-10 and in Fig.1G : rho=0.72 P< 10e-10.

Experiment 2 (hydrophilic vs. hydrophobic treated glass surfaces):

Hydrophobic surface: Dataset consisted of 7 surface sections of 1.6 mm × 1.6 mm (FoVs) from 4 different wells, with a total of 1087 droplets (1069 with at least one cell). Hydrophilic surface: Dataset consisted of 7 surface sections of 1.6 mm × 1.6 mm (FoVs) from 4 different wells, with a total of 2125 droplets (1449 with at least one cell). Data smoothing lines were done using 'smooth'(X,Y,r,'loess') with r=0.5 Fig. 2E r=0.3 Fig 2G. Spearman rank correlation coefficients in Fig.2E : hydrophobic - rho=0.14 P< 0.0294; hydrophilic - rho=0.40 P< 10e-10 (for drop area > $10^3$) and in Fig.2G: hydrophobic - rho=0.385 P= 3.4412e-10; hydrophilic - rho=0.466 P< 1.69e-29 (for drop area > $10^3$). Statistical tests in Fig. 2H were based on Welch two-sample t-test, yielded the following results at confidence level of 0.01: (1) h=1; P=0.0058; tstat: 3.3470; df: 12 ; sd: 0.0601; (2) h=0; P= 0.4103; tstat: 0.8532; df: 12; sd: 64.2169 (3) h=1; P= 1.3471e-04; tstat: 5.5071; df: 12; sd: 6.3414e-04

### Mechanistic model

Density-based mechanistic model:

A naïve mechanistic model assumes that the number of plasmid transfer events is a multiplication of the densities of donor and acceptor cells in each droplet and the droplet area, and some factor k.

$T_e = k\,(D_d \cdot D_r \cdot A)$

Where:

$D_d$ is the donor cells density (units: $1/\mu m^2$)

Dr is the recipient cell density (units: $1/\mu m^2$)

A is droplet area (units: $\mu m^2$)

k is a constant (units: $1/\mu m^2$)

To estimate k we used a simple regression model Y ~ kX where Y is the number of transfer events per droplet ($T_e$) and X equals $D_d \cdot D_r \cdot A$. Assuming p is the fraction of donors of all cells, and the recipients fraction is (1-p), and that D=C/P, we get: $T_e = k\,(D_d \cdot D_r \cdot A) = k\,[\,(p \cdot C/A) \cdot (1-p)(C/A) \cdot A]$; where C is the total number of cells (donor and recipient) in the droplet.

***Experiment 1 (untreated glass)***

We first estimated k (see section 1 below and Fig. S2) based on our data, and then replaced C based on our empirical model of cell number as a function of droplet area (see section 2 and Fig. 1C). For comparison, we also computed a best fit model (This model is based on fitting to the data, not a mechanistic model) of $T_e$ as a function of A (see section 3)

Overall $T_e$ model development:

$T_e = k\,(D_d \cdot D_r \cdot A) = k\,[p\,C/A \cdot (1-p)\,C/A \cdot A) = k[\,(p\,(\beta 1 A^{\alpha 1})/A) \cdot ((1-p)\,(\beta 1 A^{\alpha 1})/A) \cdot A]$
$= k[p(1-p)] \cdot \beta 1^2 \cdot A^{(2\,\alpha 1-1)} = 0.136 \cdot [0.24 \cdot 0.76] \cdot 0.0038^2 \cdot A^{(2.62-1)} = 3.58 \cdot 10^{-7}\,A^{1.62}$

1) *Estimating k:*

Only droplets where the average bin value crossed the threshold to show a positive number of transfers where used to calculate the slope (see Fig. S3).

Coefficients were estimated using 'fitnlm' function    Y ~ kX

Estimated Coefficients:

| | Estimate | SE | tStat | pValue |
|---|---|---|---|---|
| | ________ | _________ | ______ | __________ |
| k | 0.13619 | 0.0044114 | 30.872 | 2.4881e-60 |

Number of observations: 126, Error degrees of freedom: 125

Root Mean Squared Error: 14.9

R-Squared: 0.869,  Adjusted R-Squared 0.869

F-statistic vs. zero model: 953, p-value = 2.49e-60

*2) Cell numbers – to - drop-area model:*

Drops without any cells were removed from the analysis. Power function coefficients were estimated using 'fitnlm' function to a simple power function of the form: $Y=\beta X^{\alpha}$.
Estimated Coefficients:

| | Estimate | SE | tStat | pValue |
|---|---|---|---|---|
| β | 0.003844 | 0.0010917 | 3.521 | 0.00048631 |
| α | 1.3093 | 0.02431 | 53.859 | 2.4993e-172 |

Number of observations: 355, Error degrees of freedom: 353, Root Mean Squared Error: 1.42e+03, R-Squared: 0.896, Adjusted R-Squared 0.896, F-statistic vs. zero model: 1.86e+03, p-value = 3.12e-188

*3) Best fitted power law model based only on data:*

Estimated Coefficients:

| | Estimate | SE | tStat | pValue |
|---|---|---|---|---|
| b1 | 1.0876e-09 | 2.4986e-11 | 43.53 | 7.5508e-148 |
| b2 | 2.1293 | 3.3129e-19 | 6.4273e+18 | 0 |

Number of observations: 372, Error degrees of freedom: 371
Root Mean Squared Error: 10.3
R-Squared: 0.829, Adjusted R-Squared 0.829
F-statistic vs. zero model: 1.89e+03, p-value = 7.55e-148

***Experiment 2 (modified glass)***

Overall $T_e$ model:

$$T_e = k\,(D_d \cdot D_r \cdot A) = k\,[p\,C/A \cdot (1-p)\,C/A \cdot A) = k[\,(p\,(\beta1 A^{\alpha1})/A) \cdot ((1-p)\,(\beta1 A^{\alpha1})/A)\cdot A] = k[p(1-p)] \cdot \beta1^2 \cdot A^{(2\alpha1-1)}$$

*Treatment 1 - Hydrophobic:*

$$T_e = k[p(1-p)] \cdot \beta1^2 \cdot A^{(2\alpha1-1)} = 0.035\cdot[0.5\cdot0.5] \cdot 0.023^2 \cdot A^{(2.56-1)} = 4.62 \cdot 10^{-6} A^{1.56}$$

*Treatment 2 - Hydrophobic:*

$$T_e = k[p(1-p)] \cdot \beta1^2 \cdot A^{(2\alpha1-1)} = 0.036\cdot[0.5\cdot0.5] \cdot 0.040^2 \cdot A^{(2.32-1)} = 1.44 \cdot 10^{-5} A^{1.32}$$

1) *Estimating k:*
Only droplets where the average bin value crossed the threshold to show a positive number of transfers where used to calculate the slope (see Fig. S5).
Regression model:

$Y \sim kX$

*Treatment 1 - Hydrophobic:*

Estimated Coefficients:

| | Estimate | SE | tStat | pValue |
|---|---|---|---|---|
| k | 0.035182 | 0.0025019 | 14.062 | 8.0262e-27 |

Number of observations: 117, Error degrees of freedom: 116
Root Mean Squared Error: 12.7
R-Squared: 0.557, Adjusted R-Squared 0.557
F-statistic vs. zero model: 198, p-value = 8.03e-27

*Treatment 2 - Hydrophilic:*
Estimated Coefficients:

| | Estimate | SE | tStat | pValue |
|---|---|---|---|---|
| k | 0.036097 | 0.0013806 | 26.145 | 1.2147e-59 |

Number of observations: 161, Error degrees of freedom: 160
Root Mean Squared Error: 4.26
R-Squared: 0.738, Adjusted R-Squared 0.738
F-statistic vs. zero model: 684, p-value = 1.21e-59

*2) Cell numbers – to - drop-area model:*
Drops without any cells were removed from the analysis. Power function coefficients were estimated using 'fitnlm' function to a simple power function of the form: $Y=\beta X^{\alpha}$.
Estimated Coefficients:

*Treatment 1 - Hydrophobic:*
Estimated Coefficients:

| | Estimate | SE | tStat | pValue |
|---|---|---|---|---|
| β | 0.022876 | 0.0044684 | 5.1196 | 3.8279e-07 |
| α | 1.2815 | 0.019358 | 66.198 | 0 |

Number of observations: 812, Error degrees of freedom: 810
Root Mean Squared Error: 456
R-Squared: 0.896, Adjusted R-Squared 0.896
F-statistic vs. zero model: 3.79e+03, p-value = 0

*Treatment 2 - Hydrophilic:*
Estimated Coefficients:

| | Estimate | SE | tStat | pValue |
|---|---|---|---|---|
| β | 0.04042 | 0.0043457 | 9.3012 | 5.0394e-20 |
| α | 1.1632 | 0.010485 | 110.93 | 0 |

Number of observations: 1430, Error degrees of freedom: 1428
Root Mean Squared Error: 360
R-Squared: 0.909, Adjusted R-Squared 0.909
F-statistic vs. zero model: 7.92e+03, p-value = 0

3) Best fitted power law model based on data:

*Treatment 1 - Hydrophobic:*

Estimated Coefficients:

| | Estimate | SE | tStat | pValue |
|---|---|---|---|---|
| b1 | 8.6092e-06 | 3.1009e-06 | 2.7764 | 0.0055935 |
| b2 | 1.5385 | 0.035257 | 43.636 | 5.106e-239 |

Number of observations: 1063, Error degrees of freedom: 1061
Root Mean Squared Error: 3.11
R-Squared: 0.795, Adjusted R-Squared 0.795
F-statistic vs. zero model: 2.12e+03, p-value = 0

*Treatment 2 - Hydrophilic:*

Best fitted power law model based on data:
Estimated Coefficients:

| | Estimate | SE | tStat | pValue |
|---|---|---|---|---|
| b1 | 6.7335e-06 | 1.8278e-06 | 3.6839 | 0.00023781 |
| b2 | 1.4173 | 0.025878 | 54.77 | 0 |

Number of observations: 1515, Error degrees of freedom: 1513
Root Mean Squared Error: 1.75
R-Squared: 0.692, Adjusted R-Squared 0.692
F-statistic vs. zero model: 1.81e+03, p-value = 0

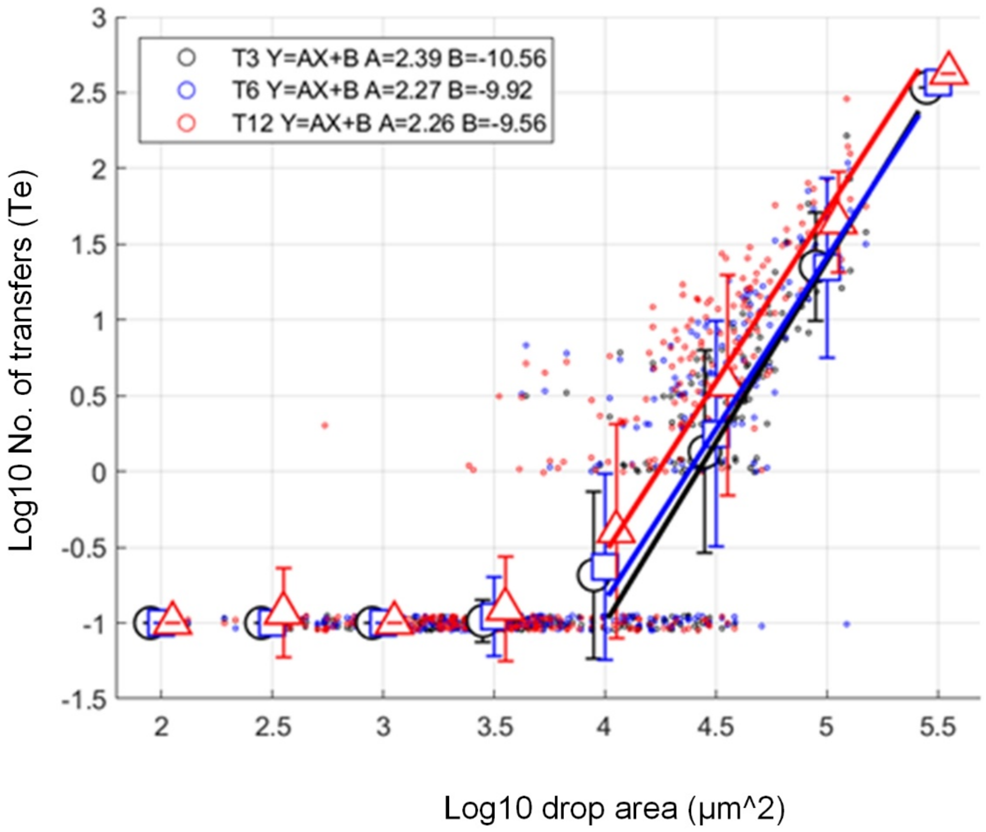


**Fig. S1.** Plasmid transfer event ($T_e$) at the three time points (3 hr [black circle], 6 hr [blue square], 12 hr [red triangle] ) follow the same pattern. Similar exponent values of (≈2.3) of the power function were observed between time points. These data indicates that the plasmid transfer events show similar dynamic pattern along these time points and across the whole range of droplet areas.

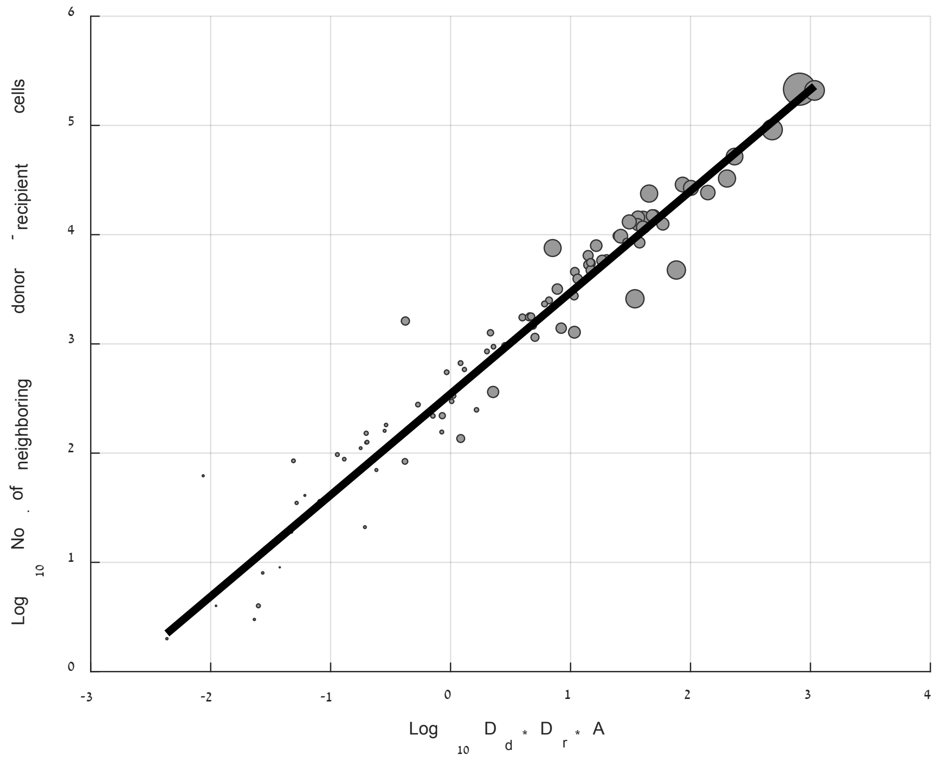


**Fig. S2. The number of neighboring donor and recipient cells within in droplet vs. mechanistic density per-droplet population-based model.** Data is shown for a subset of 106 droplets from experiment 1 (untreated glass). In each droplet the entity of each cell was determined (donor or recipient) as well as the exact location on the surface. The number of cells within a neighborhood of 5 µm was computed for each donor cell (No. of neighbors). The X-axis describe the naïve 'population-based' mechanistic model that we evaluated in the main text. Drops without any cells were removed from the analysis. Power function coefficients were estimated using 'fitnlm' function to a simple power function of the form: $Y=\beta X^{\alpha}$. This yielded the following fit: $Y=349 \cdot X^{0.93}$ with an $R^2$ of 0.97. This analysis shows that the density-based model provides a good approximation for the number of donor-recipient pairs in close proximity.

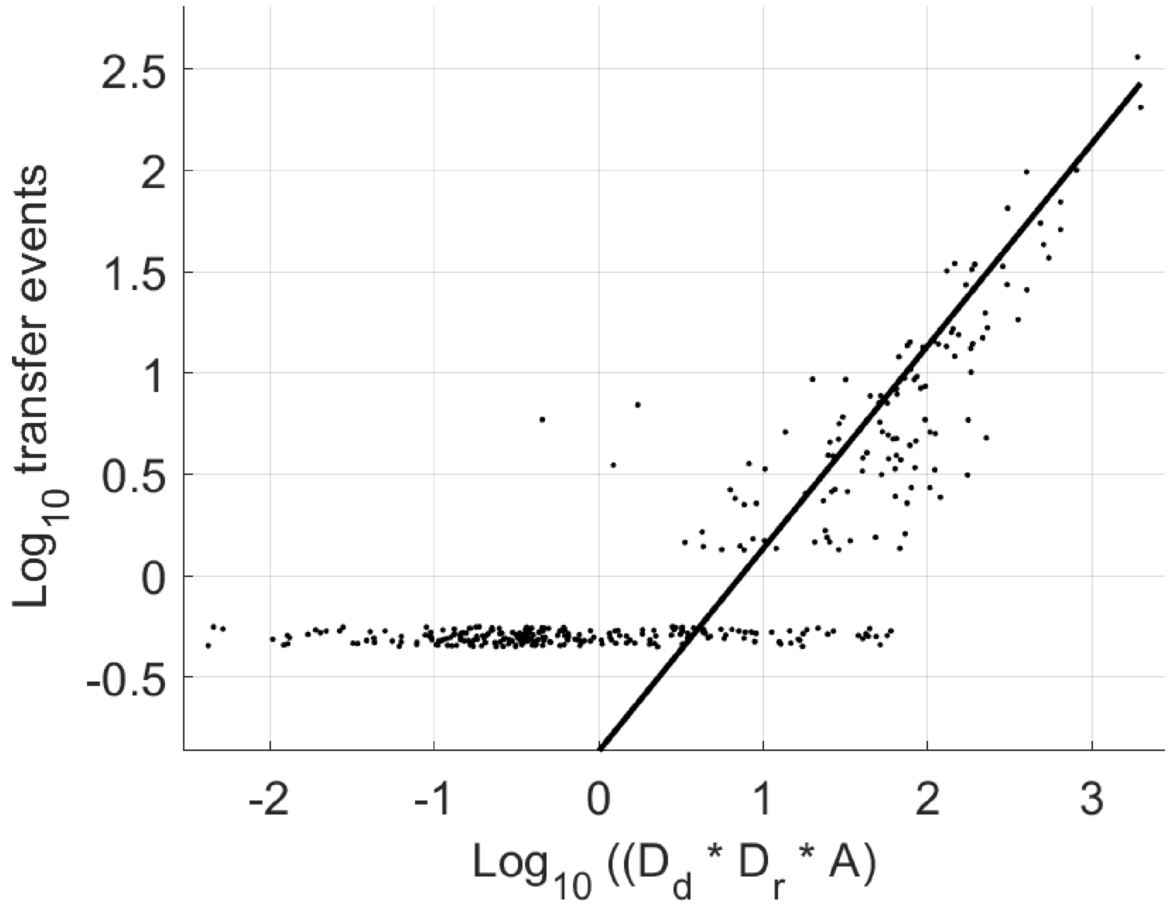


**Fig. S3.** Mechanistic Density-based model of Transfer events in experiment 1 (t= 6 hr). Axis are in log-log scales. Straight line describes fitted model based on ‘fitnlm’ Matlab function to Y=kX. k=0.136±0.004 (SE). Black dots represent experimental results.

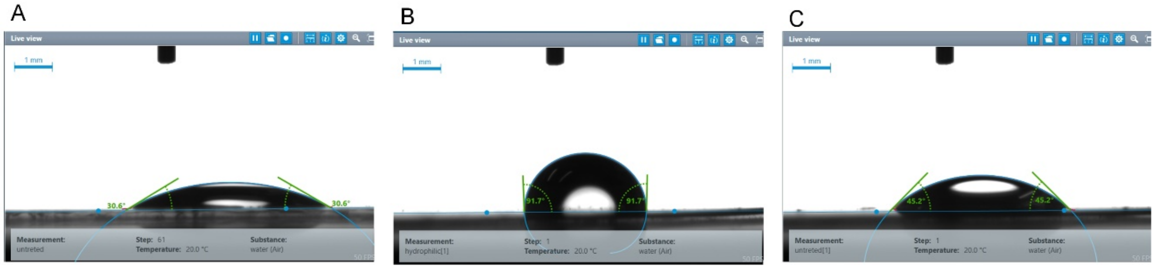


**Fig. S4. Contact angle measurements on glass and modified surfaces.** To mimic experimental conditions 11 µL drops of M9 medium supplemented with 20 mM Glucose were left for 3 hr at 28°C, 100% RH. Contact angle were measured with a goniometer (EasyDrop DSA20E, KRÜSS GmbH, Hamburg, Germany). **A.** Untreated glass: 32.34° ±2.55 (mean+SD, n=6). **B.** Treatment 1 (Hydrophobic): 90.01° ±1.97 (n=3) **C.** treatment 2 (Hydrophilic): 42.15° ±4.2 (n=4).

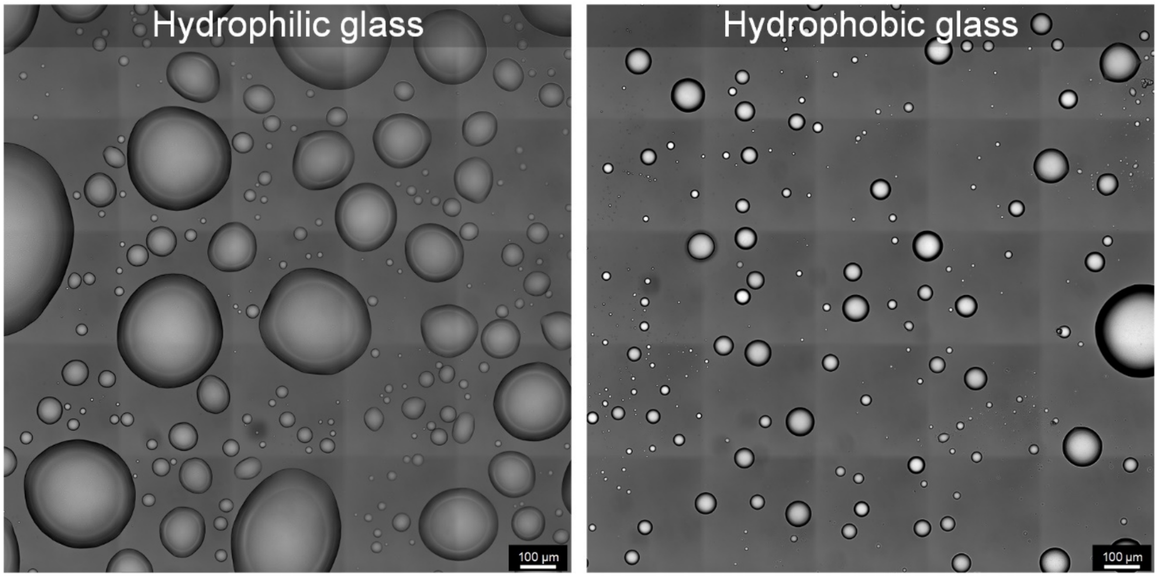


**Fig. S5.** Representative images of sprayed droplets deposited on (a) hydrophilic and (b) hydrophobic treated glass. Image constructed by stitching 5 × 5 adjacent fields of views into a single image (1.6 × 1.6 mm).

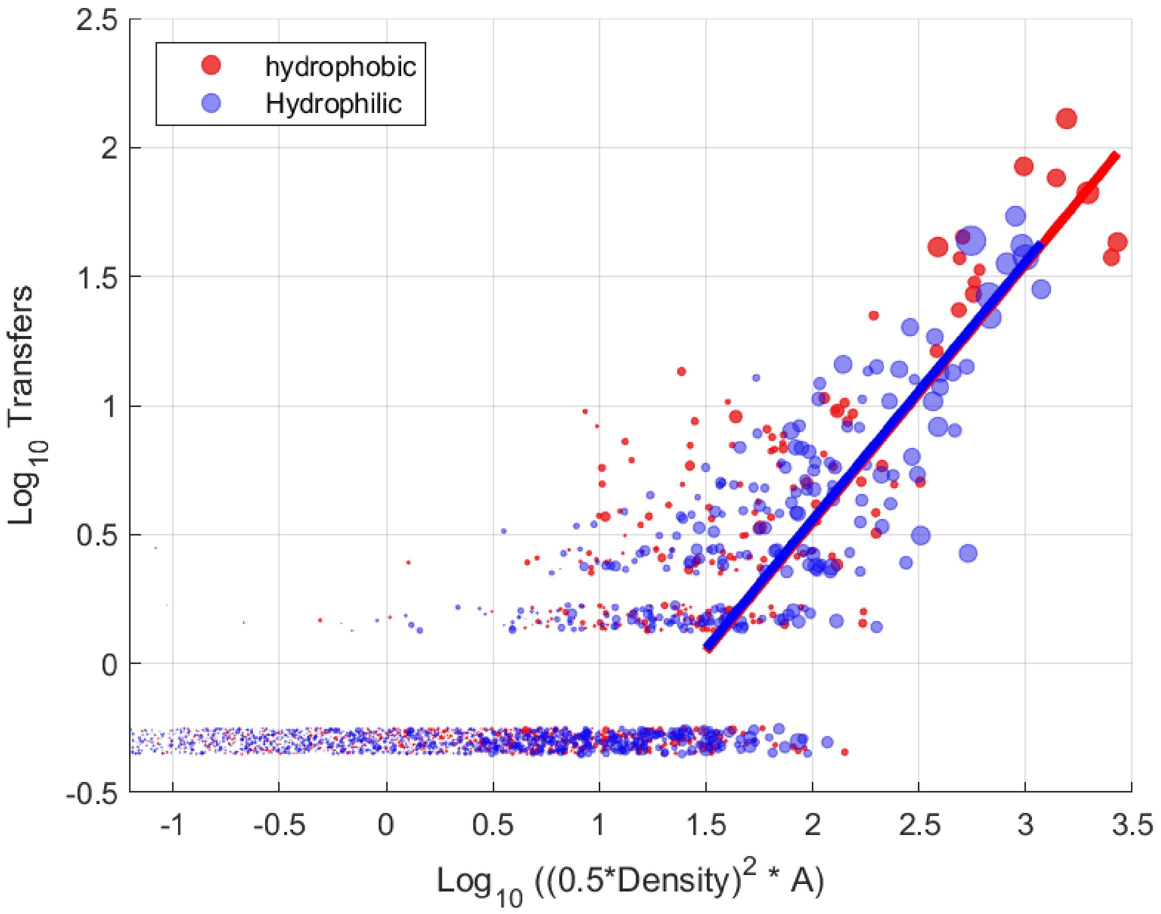


**Fig S6.** Mechanistic Density-based model of Transfer events in experiment 2: treated surfaces (t= 6 hr). Axis are in log-log scales. Straight line describes fitted model based on 'fitnlm' Matlab function to Y=kX. Red: hydrophobic surface (k1=0.035±0.002); Blue hydrophilic surface (k2=0.036±0.001). Note the fitted slope (k) is very similar (is not statistically different, 2 sample t-test).

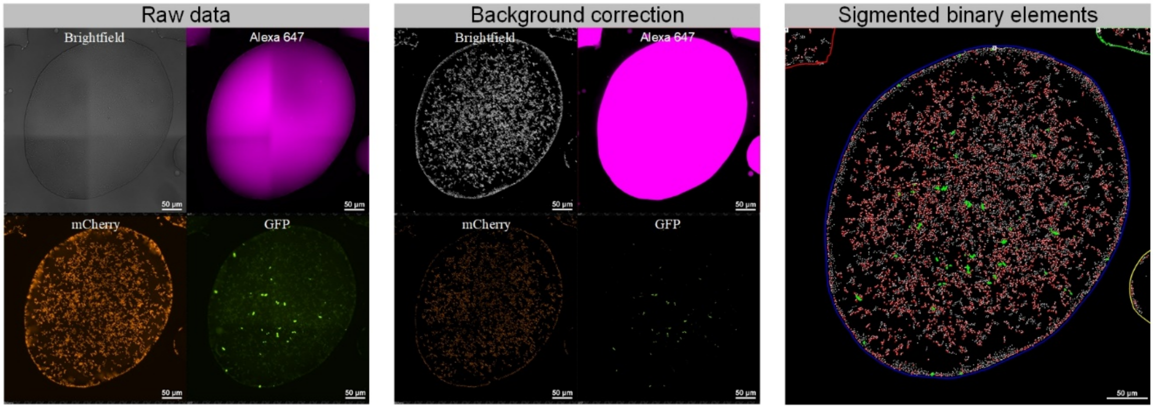


**Fig. S7.** Image processing steps. (left) Spilt channel view of raw data image. Brightfield captures the droplet and the total cells within. Alexa 647 channel captures the droplet boundaries. mCherry channel capture the donor cells. GFP channel capture trans-conjugant cells. (middle) rolling ball background correction was applied as a preprocessing step. (right) overlay of segmented elements from captured channels. Total bacterial cell in white, Donor cell in red and trans-conjugant cells in green.

| Droplet ID | No. of cells at 3hr (post inoculation) | No. of cells at 6hr | No. of cells at 12hr | No. of cells at 18hr | Log10 drop area [μm²] | Mean no. of cell divisions (per cell) |
|---|---|---|---|---|---|---|
| 1 | 5554 | 6590 | 7836 | 7951 | 4.80 | 0.72 |
| 2 | 226 | 313 | 358 | 362 | 3.76 | 0.80 |
| 3 | 321 | 467 | 607 | 609 | 3.97 | 0.95 |
| 4 | 15214 | 17555 | 21430 | 20702 | 5.09 | 0.68 |
| 5 | 2930 | 3746 | 4963 | 4789 | 4.62 | 0.81 |
| 6 | 878 | 1239 | 1507 | 1473 | 4.27 | 0.84 |
| 7 | 1833 | 2465 | 3134 | 3040 | 4.53 | 0.83 |

**Table S1.** number of cells in individual droplets over time, showing less than one cell division (on average) between 3 hr and 18 hr.
